# Ranking Single Fluorescent Protein Based Calcium Biosensor Performance by Molecular Dynamics Simulations

**DOI:** 10.1101/2024.07.29.605619

**Authors:** Melike Berksoz, Canan Atilgan

## Abstract

Genetically Encoded Fluorescent Biosensors (GEFBs) have become indispensable tools for visualizing biological processes in vivo. A typical GEFB is composed of a sensory domain (SD) which undergoes a conformational change upon ligand binding and a genetically fused fluorescent protein (FP). Ligand binding in the SD allosterically modulates the chromophore environment and changes its spectral properties. Single fluorescent (FP)-based biosensors, a subclass of GEFBs, offer a simple experimental setup; they are easy to produce in living cells, structurally stable and simple due to their single-wavelength operation. However, they pose a significant challenge for structure optimization, especially concerning the length and residue content of linkers between the FP and SD which affect how well the chromophore responds to conformational change in the SD. In this work, we use classical all-atom molecular dynamics simulations to analyze the dynamic properties of a series of calmodulin-based calcium biosensors, all with different FP-SD interaction interfaces and varying degrees of calcium binding dependent fluorescence change. Our results indicate that biosensor performance can be predicted based on distribution of water molecules around the chromophore and shifts in hydrogen bond occupancies between the ligand-bound and ligand-free sensor structures.

## 1. Introduction

Genetically encoded fluorescent biosensors (GEFBs) are widely used as molecular live imaging tools. Virtually any analyte can be traced in the transgenic cells expressing the fluorescent sensor which binds selectively to the analyte of interest. Two main classes of GEFBs are Förster Resonance Energy Transfer (FRET)-based sensors and single fluorescent protein (FP)-based sensors. FRET sensors are composed of two FPs and a sensing protein which undergoes a large enough conformational change upon ligand binding that effectively alters the distance between the two FPs, allowing for energy transfer between their fluorophores detectable as a change in fluorescence intensity.^1^ Single FP biosensors are constructed by fusion of a single FP and a sensory domain. Circular permutation of the FP allows for insertion of a sensing protein in proximity of the chromophore. Fluorescence is modulated by ligand binding/unbinding and the subsequent changes in the hydrogen bonding pattern around the chromophore.^23^ Excitation with light of suitable wavelength leads to acidification of phenolic hydrogen of the chromophore which is transferred via a network of hydrogen bonds to a nearby glutamate residue. The remaining anionic phenolate moiety displays higher fluorescence than the neutral form.^45^ Therefore, a shift in the hydrogen bond pattern around the chromophore because of a conformational change directly affects the fluorescence output of the sensor.

Genetically encoded calcium indicators (GECIs) are the earliest and most widely studied single FP biosensors given the important role of calcium in neural activity and as a secondary messenger in intracellular signaling (Figure 1). As of 2024, the largest group of fluorescent biosensors is by far the calcium sensors.6 Two main strategies are used in the design of GECIs; (i) a circularly permuted FP is fused to calmodulin (CaM) on its C terminus and to a CaM binding peptide on its N terminus, (Figure 1G) (ii) CaM-peptide moiety is inserted into the FP sequence (Figure 1H). The α helical peptide hydrophobically interacts with CaM and ‘holds’ the two domains together.^7^ Inside the FP’s *β* barrel, the chromophore responsible for the spectral properties exists in an equilibrium of neutral and anionic forms.^8^ Ca^2+^ binding triggers a conformational change in CaM which translates to a shift in the pKa of the chromophore and moves the equilibrium towards the anionic form.3 In some cases, the extinction coefficient and quantum yield may also be altered as a result.^9^ For most GECIs, the anionic chromophore is the brighter form, therefore Ca^2+^ binding causes a detectable increase in fluorescence intensity. Structural studies of GCaMP2 showed that in the calcium saturated form, the two domains of CaM wrap around the M13 peptide, effectively blocking the solvent access to circular permutation (cp) site which helps keep the chromophore in its anionic-fluorescent state.^10^ In fact, reduced solvent access to the chromophore may be common feature of biosensors in the high fluorescent state.^11^

**Figure 1.**
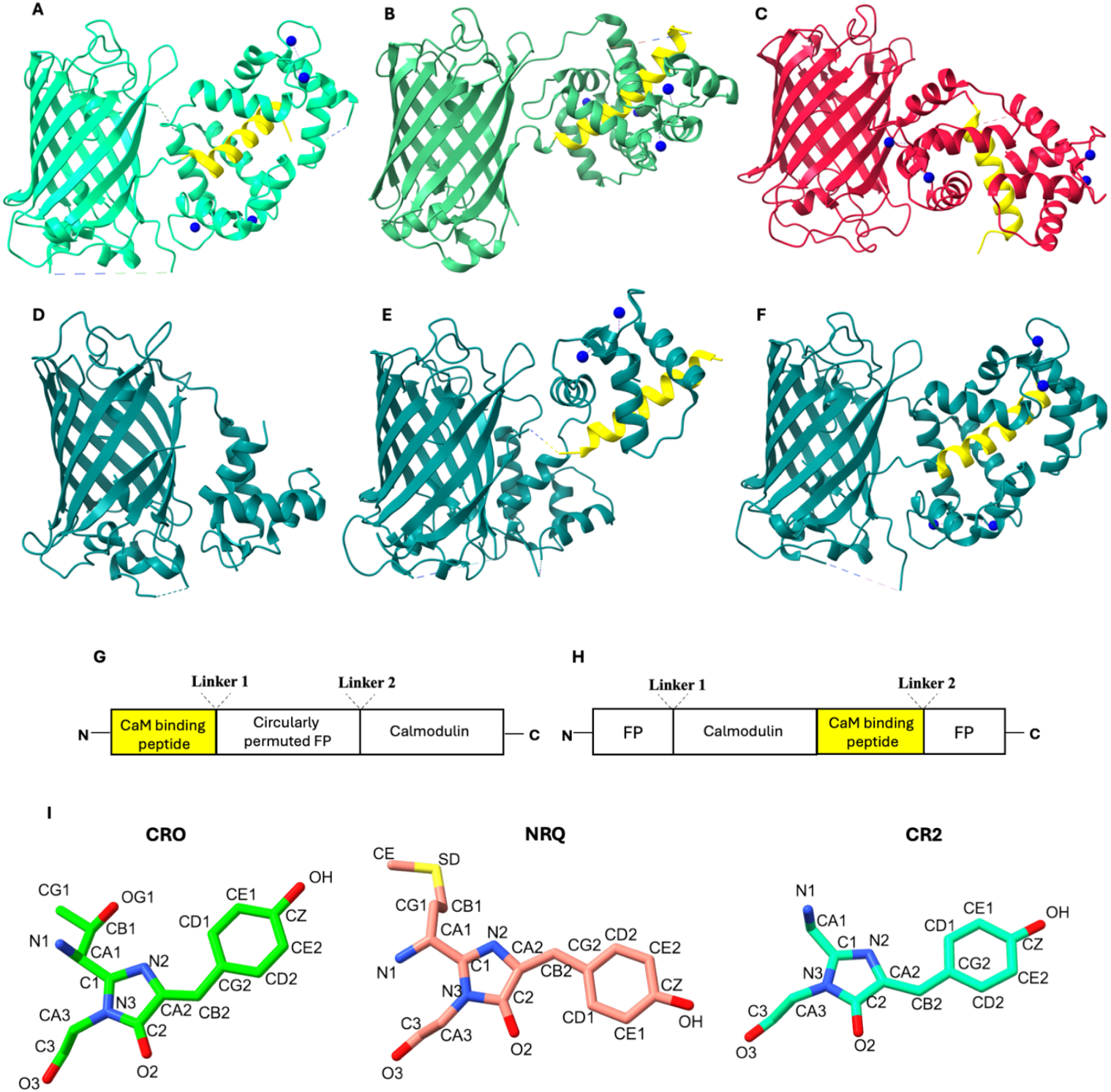
**A**. Four Ca2+ bound jGCaMP8 (PDB: 7ST4) **B**. Four Ca2+ bound NCaMP7 (PDB:6XW2) **C**. Four Ca2+ bound RCaMP1h (PDB: 3U0K) **D**. Calcium-free GCaMP2 (PDB:3EKJ) **E**. Two Ca2+ bound GCaMP2 (PDB:3O77) **F**. Four Ca2+ bound GCaMP2 (PDB:3EVR). **G**. Schematic sequence of jGCaMP8, RCaMP1h and GCaMP2 **H**. Schematic sequence of NCaMP7. **I**. Chromophore types. CRO, CR2 and NRQ. Atom types are taken from CHARMM36 force field topologies.

Two-calcium-bound GCaMP2 has an extended structure where the N and C terminal domains of CaM are distal to each other and M13 interacts only with the N terminal domain carrying the two calciums (Figure 1E). To date, there is only one published calcium-free structure of GCaMP2 and in that structure only the FP and the N terminus domain of CaM are resolved while M13 peptide and the C terminal domain are missing (Figure 1D). *Holo* structure of other GCaMP-like GECIs reveal a similar organization although the positioning of the CaM-peptide relative to the FP domain differs depending on the circular permutation strategy (Figure 1A-1C).

Biosensor performance is generally ranked based on a few biophysical features; ligand binding affinity *K*_*d*_, response rate, extinction coefficient, quantum yield and ligand binding dependent change in fluorescence signal amplitude Δ*F/F*.^12–14^ Among these, ΔF/F may be a suitable property to deduce from molecular dynamics (MD) simulations where ligand binding dependent conformational change can be sampled within reachable timescales.^15^ High Δ*F/F* might mean one of two things; there may be a large conformational change upon ligand binding in the SD which greatly changes the local environment and the pKa of the chromophore; alternatively, a few key residues at the interface between the SD and FP may be especially effective in communicating the conformational change to the chromophore. When comparing a series of sensors with the same SD, as in CaM-based calcium biosensors, the latter option seems to be decisive for their varying degree of Δ*F/F* values since the level of ligand-dependent conformational change is the same in all of them. To correlate the protein dynamics obtained from MD with fluorescence, structural requirements for the chromophore should be defined. Crystal structures of various biosensors revealed a few key properties of the ON state sensors; (i) anionic chromophore is stabilized by a direct or a water mediated hydrogen bond with a nearby donor residue located on FP, linker or sensory domain, (ii) the two rings of the chromophore are coplanar as in parental FPs, and (iii) opening at the cp site where the chromophore protrudes is partially occluded by the SD which reduces the solvent exposure and help preserve the anionic high fluorescent state.^16–20^

In this work, we analyze a series of GECIs with varying ligand dependent Δ*F/F* values with the motivation to reveal the allosteric mechanism of conformational change and its effect on chromophore environment with classical all-atom MD simulations. We have chosen green GECIs, GCaMP2^10^, NCaMP7 ^21^ and jGCaMP8 ^22^ and red GECI RCaMP1h^23^. Differences in parental FPs, chromophore types, calcium binding dependent Δ*F/F* values, CaM binding peptides and circular permutation strategies have motivated us to choose these sensors for a detailed investigation to propose a unified working mechanism that may also be predictive for new designs. Calcium-saturated crystal structures of all sensors are available in the PDB (Figures 1A-D). GCaMP2 is the earliest designed sensor in our set. It is an improved version of the original GCaMP, the first single FP based Ca^2+^ indicator and is derived from circularly permuted EGFP (Figure 1D-1F). RCaMP1h is derived from fusion of circularly permuted mRuby and CaM-M13 (Figure 1C). In NCaMP7, CaM-M13 moiety is inserted into the mNeongreen (Figure 1B). This type of design was demonstrated to be advantageous in terms of calcium sensitivity and dynamic range.^21,24^ The most recent and the superior one among the four sensors is jGCaMP8. Initial variant ‘JGCaMP8.410.80’, whose crystal structure was resolved, has a Δ*F/F* of 75.^22^ It is composed of cpGFP, CaM and the CaM-binding peptide from endothelial nitric oxide synthase (ENOSP) which has a considerably different sequence than the M13 peptide (Figure S1).

As part of design of single FP-based sensors, opening created at the cp site on FPs exposes the chromophore to nearby ionizable residues as well as solvent molecules which may protonate the anionic phenyl group and quench the fluorescence. We test the degree of exposure to solvent molecules by measuring Solvent Accessible Surface Area (SASA) of the chromophore and radial distribution of water molecules around it. Sensors have different chromophores depending on the parental FPs: ‘CRO’, ‘CR2’ and ‘NRQ’ are three kinds of chromophores found in our selection which are formed by three-residue sequences ‘TYG’, ‘GYG’ and ‘MYG’ respectively (Table 1 and Figure 1I). Along with the sensors, we analyzed four parental intact FPs: GFP, mRuby and mNeonGreen bearing CRO, NRQ and CR2 respectively. We investigate whether a higher Δ*F/F* correlates with a sensor structure where the opening created by circular permutation is occluded by the CaM-peptide domain to a greater degree. This is expected to lead to a similar environment for the chromophore to that of its parental FP. We quantify the environment of the chromophore via its solvent accessibility and the radial distribution of water molecules around it. We further propose a mechanistic explanation to the regulation of fluorescence state based on shifts in occupancies of hydrogen bonds throughout the protein. We focus on GCaMP2 and jGCaMP8 as the two extremes of low and high-performance sensors throughout the main text, while data on NCaMP7 and RCaMP1h are largely provided in Supplementary Information.

**Table 1.** Calcium biosensors and parental FPs used in this study.

| Sensor/FP | PDB code | Glutamate with upshifted pKa * | Chromophore type | Parental FP | $\Delta F/F$ |
| --- | --- | --- | --- | --- | --- |
| *GFP* | 1EMA | E222 → 8.3 | CRO | - | - |
| *mRuby* | 3U0M | E215 → 4.4 | NRQ | - | - |
| *mNeongreen* | 5LTR | E210 → 8.3 | CR2 | - | - |
| *GCaMP2* | 3EVR | E135 → 7.4 | CRO | GFP | 4 |
| *RCaMP1h* | 3U0K | E133 (E97) → 12.5 | NRQ | mRuby | 11 |
| *NCaMP7* | 6XW2 | E45 (E35) → 11.2 | CR2 | mNeongreen | 27 |
| *jGCaMP8* | 7ST4 | E106 (E95) → 7.8 | CRO | GFP | 75 |

\*Index of glutamate residues in our MD inputs are given in parentheses in cases they are different from PDB files. Calculated pKa is given following the arrow.

## 2. Methods

### 2.1. Modelling the initial coordinates of intact FPs and *apo*/*holo* sensors

We used ColabFold to complete the missing residues and to obtain an *apo*-like model for each sensor.^25^ Each *holo* sensor was modelled by providing the *holo* crystal structures given in Table 1 as homology templates. To obtain *apo* coordinates, we provided the two-calcium-bound GCaMP2 structure in the PDB coded 3U0K as template. However, we were able to obtain a relevant *apo* model only for GCaMP2, which we named *Apo*\*, while all other ColabFold predictions of *apo* forms resembled the *holo* structures although these were not provided as template. *Apo*-like models were then obtained by manually removing the four calcium ions from the *holo* coordinates of the respective GEFBs. Three-residue sequences ‘TYG’, ‘GYG’ and ‘MYG’ were used in place of CRO, CR2 and NRQ respectively as sequence inputs because ColabFold does not recognize non-standard residues.^26^

We used sequences without the purification tags and unresolved residues; therefore, residue indices are different in AF2 models (Figure S1). Highest ranked models in terms of prediction accuracy were selected, structurally aligned with their respective PDB templates and the coordinates of the cyclized chromophores were transferred into the models. *Holo* models additionally contained four calcium atoms.

All four sensors are in high fluorescent state when calcium is bound. Therefore, we modelled all *holo* structures with an anionic chromophore and all *apo* structures with a neutral chromophore reflecting the ON and OFF states, respectively. The charge states of ionizable groups were predicted with Propka3.^27^ E222 of GFP, which is known to act as an acceptor in the light-induced proton transfer reaction, has an up-shifted pKa.^28^ E95 of jGCaMP8, E35 of NCaMP7, E97 of RCaMP1h, E135 of GCaMP2 and E210 of mNeongreen structurally align with E222 and all have upshifted pKas, thus were modelled as neutral in the ON states (Table 1 and Table S1). GFP, mRuby and mNeongreen were modelled with an anionic chromophore, reflecting the ON state (Table S1).

### 2.2. MD simulations and postprocessing

Simulation boxes were prepared with VMD and simulated with NAMD packages.^29,30^ Each protein was placed in a box of water with 150 mM KCl. At least 10 Å layer of water in each direction from any atom in the system was added, so that there is at least 20 Å padding around the protein. All atoms were modelled using CHARMM36 force field.^31^ The topologies of the neutral chromophores were used as described in CHARMM36 for residue labels ‘NRQ’ ‘CR2’ and ‘CRO’. Topologies of chromophores in deprotonated form were derived by changing the charge distribution of atoms CE1, CE2, HE1, HE2, CZ and OH based on the topology of phenoxy group in CGenFF. The rest of the chromophore remained unchanged. Atom trajectories were calculated using the Verlet algorithm with a timestep of 2 fs. Particle mesh Ewald method with a cutoff distance of 12 Å was used to calculate long-range electrostatics. Each system was subjected to minimization before running the in the NPT ensemble at 310 K and 1 atm for at least 400 ns (Table S1). Coordinates were saved every 2 ps for trajectory analysis. Occupancies of hydrogen bonds involving the chromophore atoms were calculated using VMD hydrogen bond plugin with 3.5 Å donor-acceptor distance and 30° angle criteria. Occupancies of hydrogen bonds involving only standard residues were calculated with VMD Timeline plugin and all hydrogen bonds between the atoms of two residues were merged into a single occurrence using previously published Python scripts.^32,33^ To select a homogeneous conformational population for further analysis, we carried out a cluster analysis using CPPTRAJ with k-means algorithm based on Cα root mean square deviation (RMSD).^34^ Three conformational clusters were generated and their fractions over the trajectory were plotted (Figure S2). Chromophore SASA was calculated using the Shrake-Rupley algorithm by a VMD Tcl script with a probe radius of 1.4 Å. Radial distributions of water oxygen atoms up to 10 Å distance Channels connecting the chromophore to the surface of the protein were calculated with MOLE 2.5 web interface.^35^ Representative frames obtained from MD simulations were uploaded to the webserver; channels were calculated with a bottleneck radius of 1.5 Å, bottleneck tolerance of 3 Å and maximum tunnel similarity of 0.3, with ‘Ignore HETATMs’ option off to account for the space occupied by the chromophore. Channels were visualized with ChimeraX. All the calculations were performed on the combined equilibrated trajectories of replicate runs (gray shaded regions in Figure S2). *Holo*-to-*apo* morph movies were created with ChimeraX ‘morph’ command.^36^

## 3. Results

### 3.1. MD equilibration and *holo*-to-*apo* transition upon calcium removal

We observed different level of conformational change in response to calcium removal in the CaM domains among the four sensors studied. RMSD of CaM-peptide domain between the dominant conformations of *apo* and *holo* are 3.9 Å for jGCaMP8, 2.3 Å for GCaMP2, 2.4 Å for RCaMP1h and 3.4 Å for NCaMP7. RMSD plots show a greater backbone mobility for *apo* runs in all four sensors (Figure S2). *Apo*\* run of GCaMP2 which started from the coordinates of two calcium-bound template (PDB:3O77) has even greater mobility. In this structure the two lobes of CaM are distant from each other where M13 peptide is closer to the N terminal lobe. MD run showed a deviation from this structure; we observed that the two lobes approach each other, and the CaM domain becomes more compact. This conformation becomes dominant after nearly 30 ns (pop0 in *Apo*\* plot in Figure S2). All intact FPs showed considerably lower mobility and faster equilibration than the sensors in the ON state (Figure S3).

CaM domain, which is the common sensory domain in all, contains four EF-hand motifs. Change in EF2 of jGCaMP8 (in sequential order) is displayed in Figure S4C. At this site, calcium is coordinated by one glutamate, three aspartate, one threonine and one water. Upon removal of the calcium, coordinating residues pull away from each other due to electrostatic repulsion. A similar reorganization is observed in the other three calcium binding sites; although the exact content of binding residues differs while one water molecule is present in all (data not shown). JGCaMP8 and GCaMP2 have similar positioning of cpGFP and CaM domains relative to each other and calcium removal has a similar effect in both (Figure S4A and S4F). JGCaMP8’s CRO is hydrogen bonded to Y341, mostly through a water molecule, on helix 328-341 whereas GCaMP2’s CRO makes a direct hydrogen bon to R377 on the same helix (F368-K380 in 3EVR numbering) (Figure 2A and 2D). Upon calcium removal, this helix, along with the rest of the CaM domain, pulls away from the FP domain and is slightly distorted in both GCaMP2 and jGCaMP8 (Movies S1 and S2, respectively). Furthermore, we see a progressive change (gain or loss) of interaction with residues from within the *β* barrel going from *holo* to fully *apo* state (Figure S5D). In both JGCaMP8 and GCaMP2, there is a significant reorganization of the chromophore lining residues within the *β* barrel in the *holo* to *apo* transition (Movies S1 and S2). This level of change of positions of *β* barrel residues is not observed in NCaMP7 and RCaMP1h (Movies S3 and S4).

**Figure 2.**
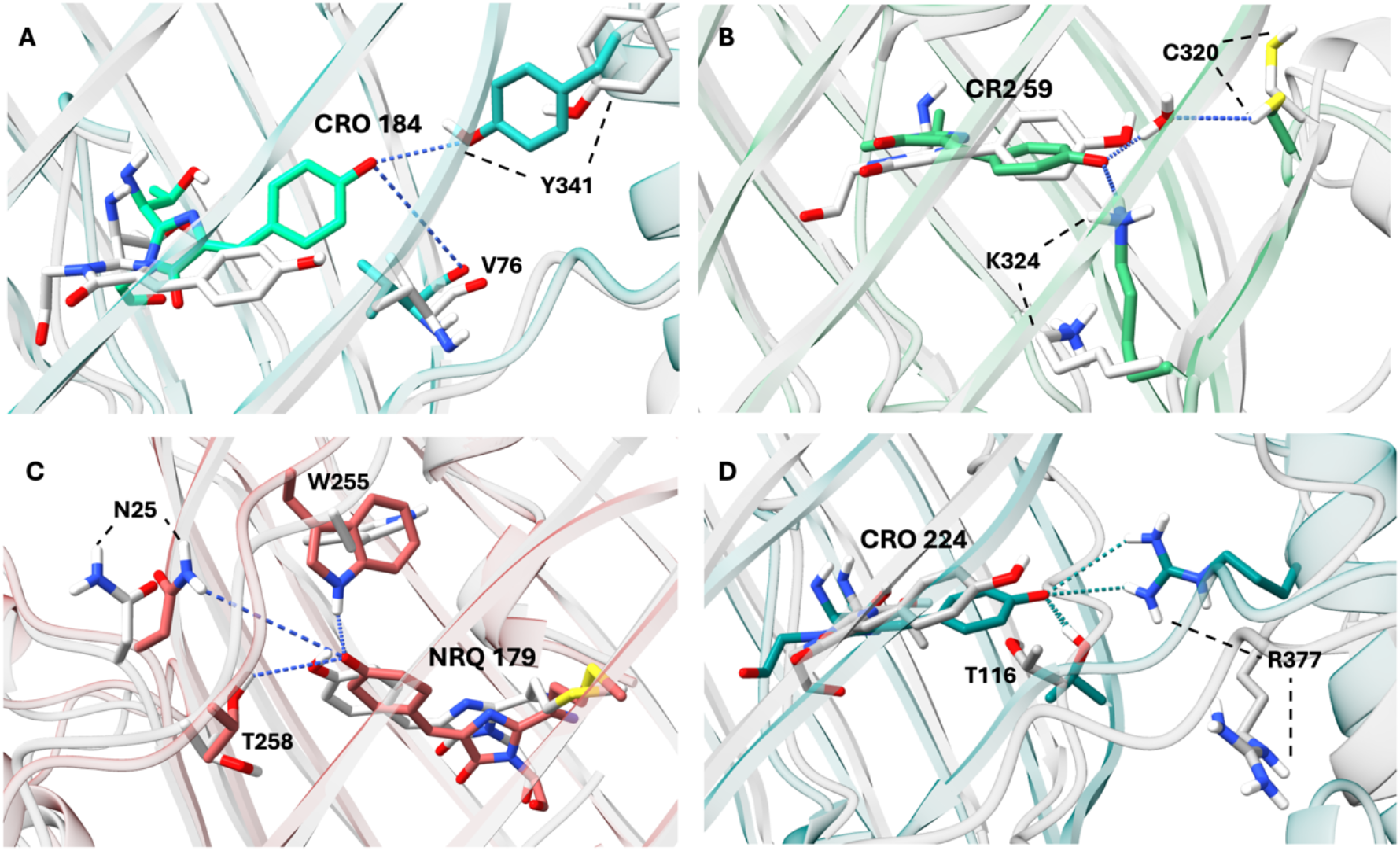
Hydrogen bond network near the chromophore in the ON and OFF state of each sensor. OFF states are colored gray and ON states are colored according to color emitted by the sensor. Images taken from representative frames of each MD run and matched with ChimeraX onto FP domains. **A**. jGCaMP8 **B**. NCaMP7 **C**. RCaMP1h **D**. GCaMP2

Although RCaMP1h has the same circular permutation strategy as GCaMP2 and JGCaMP8, the position of the M13 peptide relative to FP in the three-dimensional structure is quite different (Figure 1C). The crystal structure reveals T258 as the only hydrogen bond donor to the phenoxy oxygen of the chromophore. Our results indicate that W255 on the FP and N25 on the linker also take part in stabilizing the anionic chromophore (Figure 2C and Figure S5C), although interactions with T255 and N25 are mediated by water molecules.

In NCaMP7, intact CaM-M13 sequence is inserted into mNeongreen (Figure 1H). Because of the covalent connection to the CaM domain, M13 helix moves along with the CaM domain and changes its position relative to FP to a greater degree than other three sensors (Figure S4D). There is a clear shift in the position of hydrogen bonded residue pairs at the cp site, as well as the hydrogen bond donors of the anionic chromophore (Movie S4). The NCaMP7 chromophore has two hydrogen bond donors on FP, K324 and one gate post residue C320 (Figure 2B). High occupancy of K324-NRQ in *holo* state (80 %) relative to *apo* is probably due to the negative charge on the chromophore rather than a conformational difference (Figure S5B). The hydrogen bond with C320 is mediated by a water molecule. The distance between the chromophore phenoxy oxygen and its primary hydrogen bond donor increases when going from *holo* to *apo* state in all three sensors, except for NCaMP7 (Figure S6). CR2(OH)-C320(SH) distance in the *apo* state still allows for a water mediated hydrogen bond (Figure S6B).

### 3.2. Hydrogen bond networks distinguish the dark vs. bright states

To reveal the allosteric effect of calcium removal to the chromophore environment, we analyzed the shift in hydrogen bond occupancies over the whole structure. We listed the residue pairs with altered hydrogen bond occupancies beyond a certain threshold to discern the shifts in hydrogen bonded residue pairs. We determined the threshold for each system based on the distribution of occupancy differences between *holo* and *apo*. Similar to our previous work on a maltose biosensor, the majority of hydrogen bond occupancies remain within ± 50 % change interval when the *apo* and *holo* runs are compared.11 We found that the shift in hydrogen bond occupancies characterizes the conformational change in the CaM domain that is communicated to the chromophore environment. In fact, a distinctive feature of ON state sensors is a continuous network of hydrogen bonds from the calcium binding site to the chromophore environment. Figure 4 shows the location of hydrogen bonded residue pairs in GCaMP2 whose occupancies change by more than 50 % when going from *holo* to *apo* state. *Holo* state has considerably more hydrogen bonded pairs in calcium binding sites, as well as at the interface between the FP and CaM domain. These interactions are progressively lost when going from *holo* to *Apo*\* state (A to C) (Figure 3A, 3B and 3C and Table S2). When comparing the *Holo* and *Apo*\* states, higher number of hydrogen bonds are completely lost or gained (Figure 3D vs 3E) which is revealed by higher number of occurrences with ±100 % change. Still, manual removal of calcium ions and the resulting *apo*-like structure (Figure 3B) provides sufficient insight for changes occurring in hydrogen bond distribution, especially of residue pairs at FP-CaM interface, leading to loss of fluorescence. However, one must be cautious as the difference in hydrogen bond occupancies observed between *holo* and *apo* states would depend on how well the *apo* state is sampled starting from *holo* coordinates. For example, in case of RCaMP1h, some interactions around the linker area linger on in the *apo* state (Figure S7G vs S7H).

**Figure 3.**
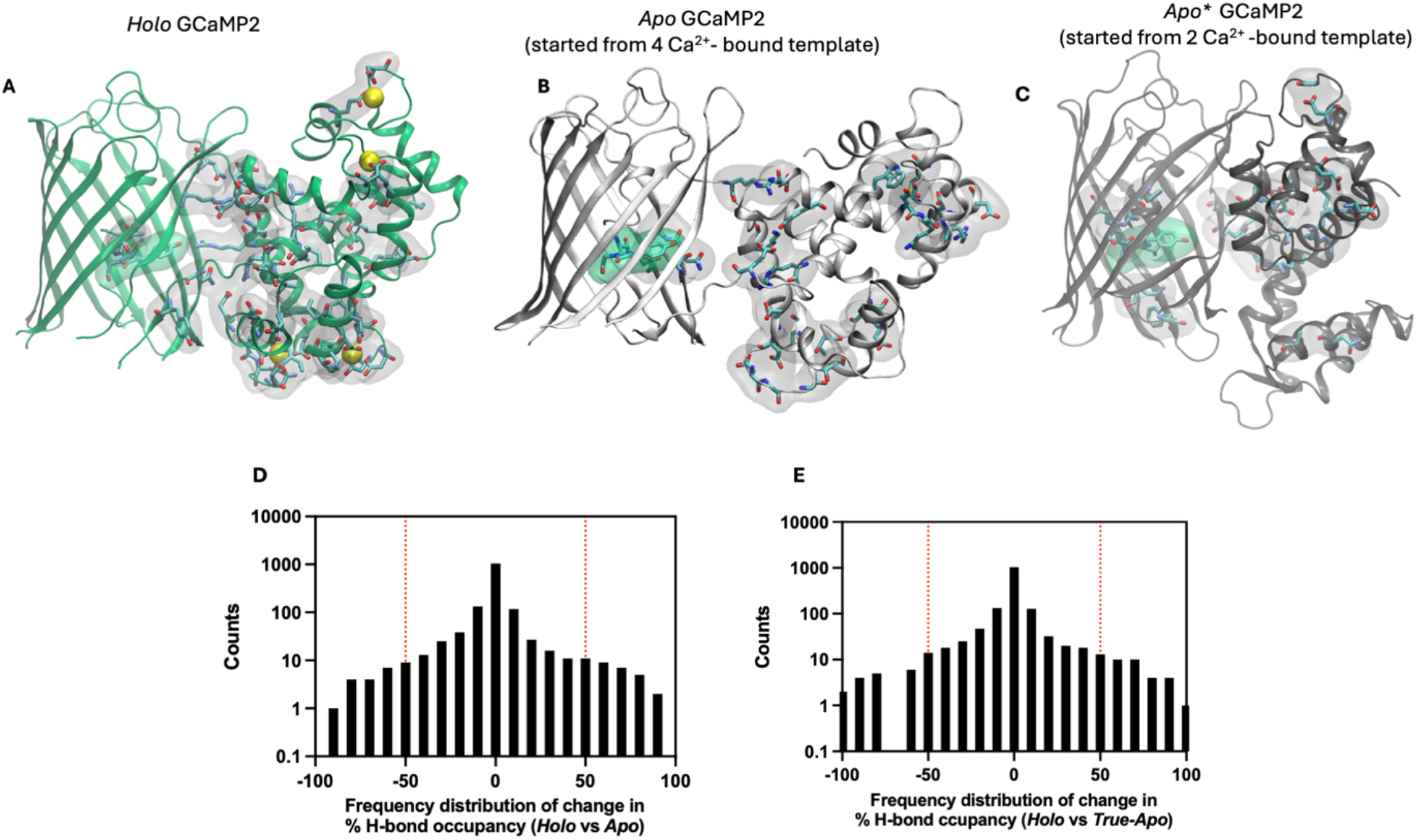
Shifts in hydrogen bond occupancies between *holo* and *apo* states of GCaMP2. Hydrogen bonded residue pairs whose occupancy increase by more than 50 % *holo* and *apo* states compared to one another are visualized as licorice and surface representations **A**. *Holo* GCaMP2 when compared to *apo* GCaMP2. **B**. *Apo* GCaMP2 when compared to *holo* GCaMP2 **C**. *Apo*\* GCaMP2 when compared to *holo* GCaMP2. Distribution of percent change in hydrogen bond occupancies **D**. between A and B **E**. between A and C. Y-axes are plotted in logarithmic scale.

**Figure 4.**
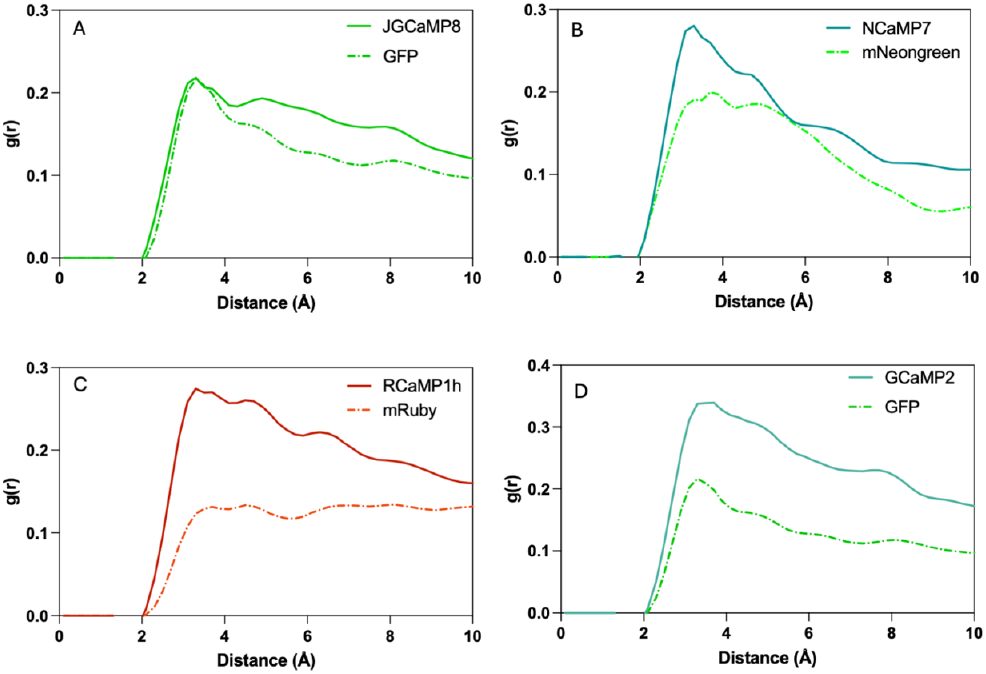
Radial distribution function of water-oxygen around chromophore in *holo* sensors and intact FPs.

### 3.4. Water number density around the chromophore is a direct indicator of fluorescence efficiency

As a measure of openness at the cp site, we examined the distribution of water molecules in the vicinity of the chromophore. Radial Distribution Function (RDF or g(r)) describes the density of a chosen group of molecules around a reference point as a function of distance between them. We measured the RDF of water molecules around the chromophore in the ON state of all sensors and their parental FPs. We chose to calculate the number of water molecules up to 10 Å as this is the minimum distance between the chromophore phenoxy oxygen, which may be protonated by water, and the CaM domain. RDF plots showed a significantly reduced number of water molecules around the chromophore in all three parental FPs compared to sensors (Figure 4). This is a plausible result since the chromophore rests within the undisrupted *β* barrel. For all systems, a common peak at around 3.3 Å points to the first layer of solvation. We a found a quantitative relation between Δ*F/F* of a sensor and the ratio of area under the g(r) curve to that of its parental FP. Ratios of integrated area up to 10 Å are JGCaMP8/GFP = 1.18, NCaMP7/mNeongreen = 1.32, RCaMP1h/mRuby = 1.56 and GCaMP2/GFP = 1.83. This ordering is the same as that of Δ*F/F* among the four sensors. This result points to a distinctive feature between sensors of low and high Δ*F/F*; high Δ*F/F* sensors seem to have a smaller number of water molecules around the chromophore normalized by their intact FPs. Structurally, we could say that the opening created at the cp site is occluded by the sensory domain to a greater extent in sensors of high Δ*F/F*.

### 3.5. The more organized solvent channels around cp sites lead to more efficient fluorescence

As a qualitative evaluation of openness around the cp site, we visualized the solvent channels connecting the chromophore to the surface using Mole 2.5 software. We used the representative frames obtained from each MD run. When comparing the *holo* states of GCaMP2 (having the smallest Δ*F/F* in our set) and jGCaMP8 (having the largest Δ*F/F* in our set), more voluminous channels are found near the GCaMP2 chromophore (Figure 5A and 5B). These channels surround the residues between the chromophore, gate post residues and residues from the CaM domain interacting with the FP domain. The bulky tyrosine residue (Y341) at this site seems to fill up this space in jGCaMP8. Additionally, the side chain of gate post residue I21 also faces inwards, while both gate post residues of GCaMP2 faces towards the solvent. We also found that the distance between Cα atoms of two gate post residues are the greatest in the case of GCaMP2 (Figure 5A, 5B and S8), even though it has a very similar overall structure to jGCaMP8 (Cα RMSD of 0.3 Å between their *holo* states). A larger distance between the gate post residues also increases the likelihood of finding a channel around the cp site. Furthermore, we found that the opening around the cp site and the number of channels increases going from *holo* to *apo* state in GCaMP2 (Figure S9). Visualizing solvent channels around the cp site could be a quick way to evaluate the degree of chromophore exposure from experimental or predicted PDB structures. However, it may be limited to differentiating sensors with very high Δ*F/F* difference, as in the case for GCaMP2 and jGCaMP8. RCaMP1h and NCaMP7 showed a similar channel profile between the two extremes (data not shown).

**Figure 5.**
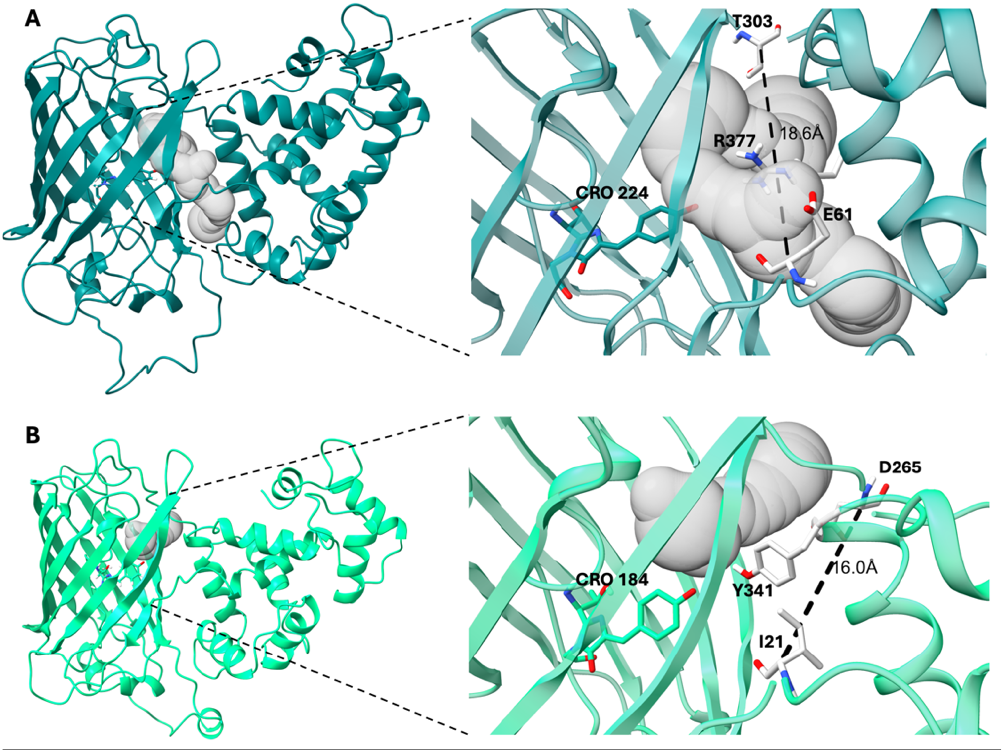
Channels connecting the surface to chromophore. **A**. *Holo* GCaMP2 **B**. *Holo* jGCAMP8. Insets shows close-up of channels and distances between gate post residues flanking the bulge.

## 4. Discussion

In this work, we used classical MD simulations to shed light on the working mechanism of a series of CaM-based single FP biosensors with varying degrees of brightness. Our simulations provided intriguing insights into the relation between biosensor dynamics and calcium binding-dependent fluorescence. One key observation was that sensors with enhanced Δ*F/F* have a limited distribution of water molecules around the chromophore. Water exposure is known to cause protonation of the phenoxy group of chromophore and quenching of the fluorescence.37 Apart from protonation, water can also alter the chromophore local environment and disrupt the hydrogen bond network within the *β* barrel.^38^ It is known that the fluorescence emission of GFP heavily depends on the hydrogen bond network and the surrounding water molecules.^4^,^39^ We found that intact FPs without circular permutation have considerably less water around the chromophore. Our analysis revealed a significant difference between the two extremes of our selection: GCaMP2 (Δ*F/F* = 4) and jGCaMP8 (Δ*F/F* > 100). The two sensors share a high structural homology despite the differences in CaM binding peptide sequences and a few mutations at the interface between FP and CaM. This suggests that MD simulations can provide guidance in evaluating randomized linker variants and reduce the experimental screening time and cost.

Our analysis of hydrogen bond occupancies between ligand-bound and ligand-free sensor structures provided additional mechanistic insights into the allosteric modulation of chromophore environment. Shifts in hydrogen bond occupancies have been shown to regulate the functioning of other proteins as well. ^32,40,41^ Our findings suggest that shift in hydrogen bond occupancy patterns upon calcium binding serves as a means to transmit conformational changes occurring in the CaM domain to the chromophore environment. The majority of hydrogen bond occupancies remains within ±50% change interval when comparing *holo* and *apo* runs. These residue pairs constitute the FP beta barrel, and they remain intact during both the *holo* and *apo* runs. Changes beyond ±50% provides us a a clue on how the conformational change effects the whole structure. Notably, a distinct characteristic of ON state sensors is the presence of a continuous network of hydrogen bonds extending from the calcium binding site to the chromophore environment. This is a common profile observed in all four sensors, irrespective of the degree of Δ*F/F* and seems to be a minimum requirement for the chromophore to function. We note that, in particular, the hydrogen bond network is disrupted near the FP-SD interface in the *apo* state. We consider this parameter as an important structural checkpoint when assessing the sensor performance. In future studies, this hypothesis could be checked by modelling failed sensor designs reported in the literature. In conclusion, our study forms the groundwork to understand structural determinants of high-performance biosensors and derive design principles from MD simulations.

## ASSOCIATED CONTENT

### Supporting Information

Supporting data on MD simulations that is not presented in the main text (.PDF)

Movie S1. *Holo*-to-*apo* transition in GCaMP2 (.mp4)

Movie S2. *Holo*-to-*apo* transition in JGCaMP8 (.mp4)

Movie S3. *Holo*-to-*apo* transition in RCaMP1h (.mp4)

Movie S4. *Holo*-to-*apo* transition in NCaMP7 (.mp4)

## AUTHOR INFORMATION

### Author Contributions

The manuscript was written through contributions of all authors. All authors have given approval to the final version of the manuscript.

## ACKNOWLEDGMENT

This work was financially supported by TUBITAK project no. 121Z329. We thank Ebru Cetin for her contribution in hydrogen bond occupancy data calculations. We thank Ali Rana Atilgan for stimulating discussions. The MD simulations in this paper were partially performed at TUBITAK ULAKBIM, High Performance and Grid Computing Center (TRUBA resources).

## ABBREVIATIONS

Cp: circular permutation
GFP: green fluorescent protein
CaM: calmodulin
SD: sensory domain
MD: molecular dynamics
RDF: radial distribution function
RMSD: root mean square deviation

