## Supplementary material for "Ranking Single Fluorescent Protein Based Calcium Biosensor Performance by Molecular Dynamics Simulations"

**Table S1. A summary of MD simulation runs**

| *System* | *Model name* | *Chromophore charge* | *MD run time* | *Replicates* |
|---|---|---|---|---|
| ***GFP*** | ON | Anionic | 400 ns | 1 |
| ***mRuby*** | ON | Anionic | 400 ns | 1 |
| ***mNeongreen*** | ON | Anionic | 400 ns | 1 |
| ***GCaMP2*** | Holo | Anionic | 600 ns | 1 |
| | Apo | Neutral | 600 ns | 1 |
| | Apo* | Neutral | 600 ns | 1 |
| ***RCaMP1h*** | Holo | Anionic | 600 ns | 2 |
| | Apo | Neutral | 600 ns | 2 |
| ***NCaMP7*** | Holo | Anionic | 600 ns | 2 |
| | Apo | Neutral | 600 ns | 2 |
| ***jGCaMP8*** | Holo | Anionic | 600 ns | 2 |
| | Apo | Neutral | 600 ns | 2 |

```
NCaMP7     MGGSHHHHHHGMASMTGGQQMGRDLYDDDDKEN--------------------------- 33
RCaMP1h    -MGSHHHHHHGMASMTGGQQMGRDLYDDDDKDLATMVDSSRRKWNKTGHAVRAIGRLSSA 59
jGCaMP8    -------------------------------MHHHHHHTRRKKTFKEVANAVKISASLM-G 29
GCaMP2     -------------------------------------DSSRRKWNKTGHAVRAIG-RL-S 21

NCaMP7     ------------------------------------------------------------ 33
RCaMP1h    INTEMMYP---ADGGLRGYTHMALKVDGGG-HLSCSFVTTYRSKKTVGNIKMPGIHYVSH 115
jGCaMP8    LKINVYIKADKQKNGIKANFHIRHNIEDGGVQLAYHY--QQNTPIGDGPVLLPDNHYLSV 87
GCaMP2     SLENVYIMADKQKNGIKANFKIRHNIEDGGVQLAYHY--QQNTPIGDGPVLLPDNHYLST 79

NCaMP7     --------------------------------LYFQGHMRSMVSKGEEENMASLPATHEL 61
RCaMP1h    RLERLEES---DNEMFVVQREHAVAKFVGL--GGGGGTGGSMNS----LIKENMR--MKV 164
jGCaMP8    QSKLSKDPNEKRDHMVLLEFVTAAGITLGMDELYKGGTGGSMVSKGEELFTGVVP--ILV 145
GCaMP2     QSKLSKDPNEKRDHMVLLEFVTAAGITLGMDELYKGGTGGSMVSKGEELFTGVVP--ILV 137

NCaMP7     HIFGSINGIDFDMVGQGTGNPNDGYEELNLKS-TMGDLQFSPWILVPHIGYGFHQYLPYP 120
RCaMP1h    VLEGSVNGHQFKCTGEGEGNPYMGTQTMRIKVIEGGPLPFAFDILATSFMYGSRTFIKYP 224
jGCaMP8    ELDGDVNGHKFSVSGEGEGDATYGKLTLKFI-CTTGKLPVPWPTLVTTLTYGVQCFSRYP 204
GCaMP2     ELDGDVNGHKFSVSGEGEGDATYGKLTLKFI-CTTGKLPVPWPTLVTTLTYGVQCFSRYP 196

NCaMP7     DGMSPFQA-AMVDGSGYQVHRTMQFEDGASLTVNYRYTYEGSHIKGEAQVEGTGFPADGP 179
RCaMP1h    KGI--PDFFKQSFPEGFTWERVTRYEDGGVITVMQDTSLEDGCLVYHAQVRGVNFPSNGA 282
jGCaMP8    DHMKQHDFFKSAMPEGYIQERTIFFKDDGNYKTRAEVKFEGDTLVNRIELKGIDFKEDGN 264
GCaMP2     DHMKQHDFFKSAMPEGYIQERTIFFKDDGNYKTRAEVKFEGDTLVNRIELKGIDFKEDGN 256

NCaMP7     VMTNSLTA--EAHDQLTEEQIAEFKEAFSLFDKDGDGTITTKELGTVMRSLGQNPTEAEL 237
RCaMP1h    VMQKKTKGWEPTRDQLTEEQIAEFKEAFSLFDKDGDGTITTKELGTVMRSLGQNPTEAEL 342
jGCaMP8    ILGHKLEY--NLPDQLTEEQIAEFKEAFSLFDKDGDGTITTKELGTVMRSLGQNPTEAEL 322
GCaMP2     ILGHKLEY--NTRDQLTEEQIAEFKEAFSLFDKDGDGTITTKELGTVMRSLGQNPTEAEL 314

NCaMP7     RVMIIEVDADGDGTLDFPEFLAMMARKMKYRDTEEEIREAFGVFDKDGNGYIGAAELRHV 297
RCaMP1h    QDMINEVDADGDGTIDFPEFLIMMARKMKDTDSEEEIREAFRVFDKDGNGYISAAELRHV 402
jGCaMP8    QDMINEVDADGDGTIDFPEFLTMMARKMKYRDTEEEIREAFGVFDKDGNGYISAAELRHV 382
GCaMP2     QDMINEVDADGNGTIDFPEFLTMMARKMKDTDSEEEIREAFRVFDKDGNGYISAAELRHV 374

NCaMP7     MTNLGEKLTDEEVGELIREADIDGDGQVNYEEFVQMMTAKGGSGGGSSSRRKWNKAGHAV 357
RCaMP1h    MTNLGEKLTDEEVDEMIREADIDGDGQVNYEEFVQMMTAK-------------------- 442
jGCaMP8    MTNLGEKLTDEEVDEMIREADIDGDGQVNYEEFVQMMTAK-------------------- 422
GCaMP2     MTNLGEKLTDEEVDEMIREADIDGDGQVNYEEFVQMMTA--------------------- 413

NCaMP7     RAIGRLSSMYFADWCVSKKTCPNDKTIVSTFKWAFITDNGKRYRSTARTTYTFAKPMAAN 417
RCaMP1h    ------------------------------------------------------------ 442
jGCaMP8    ------------------------------------------------------------ 422
GCaMP2     ------------------------------------------------------------ 413

NCaMP7     YLKNQPMYVFRKTELKHSKTELNFKEWQKAFTDVMGMDELYK   459
RCaMP1h    ------------------------------------------   442
jGCaMP8    ------------------------------------------   422
GCaMP2     ------------------------------------------   413
```

**Figure S1. Clustal Omega muliple sequence alignment of the four sensors.** Yellow and blue sequences show CaM binding peptide residues and CaM residues, respectively. Green- and salmon-colored sequences show GFP or mNeongreen and mRuby, respectively. Gray sequence are unresolved residues in the crystal structures and were not included in our initial models. Residue indices in our initial models are shifted in the following manner: **NCaMP7: n-10, RCaMP1h: n-37, jGCaMP8: n-11, GCaMP2: n**, with n being the index number in the PDB files.

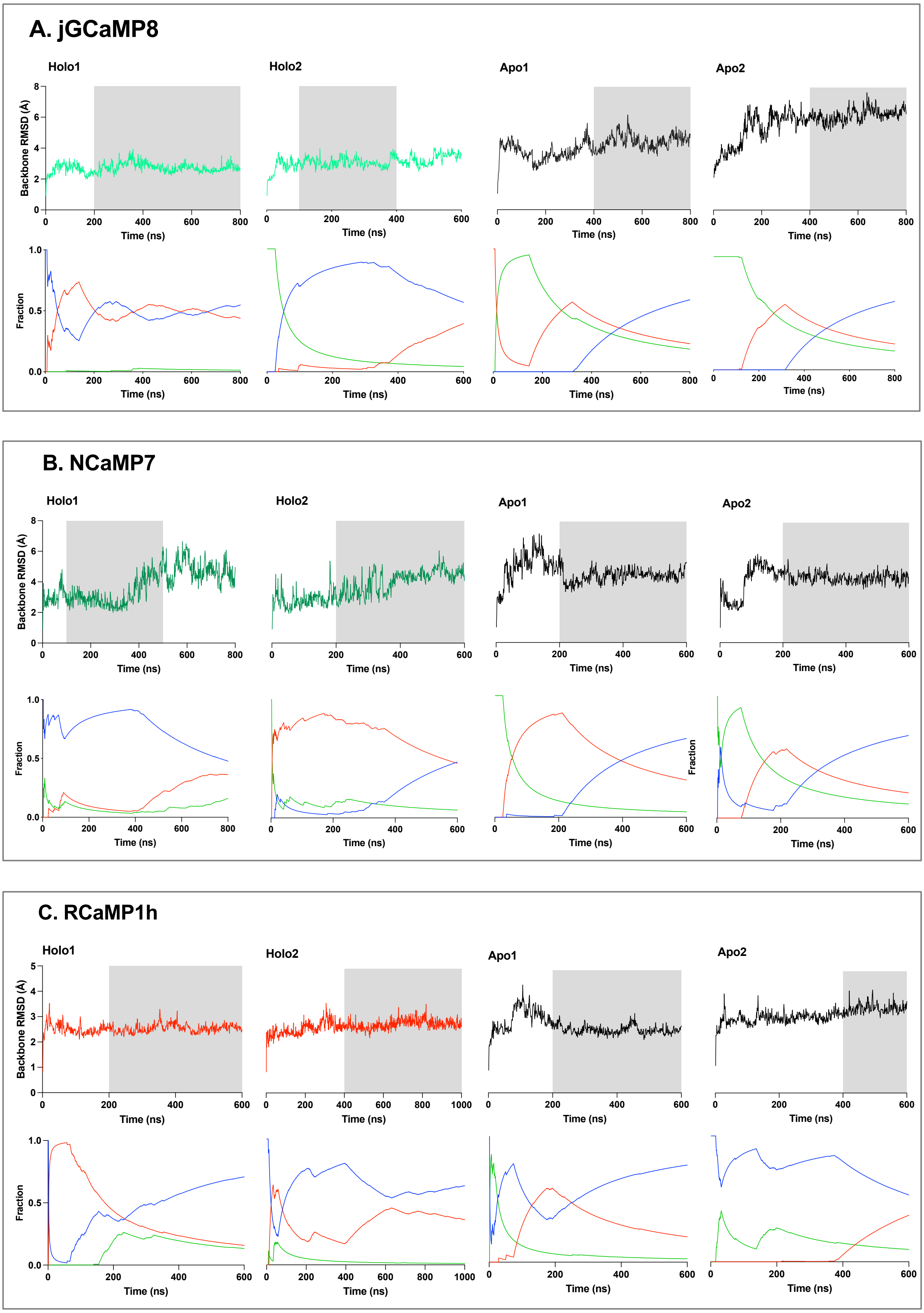
A. jGCaMP8
Holo1
Holo2
Apo1
Apo2
Backbone RMSD (Å)
Time (ns)
Fraction
B. NCaMP7
Holo1
Holo2
Apo1
Apo2
Backbone RMSD (Å)
Time (ns)
Fraction
C. RCaMP1h
Holo1
Holo2
Apo1
Apo2
Backbone RMSD (Å)
Time (ns)
Fraction

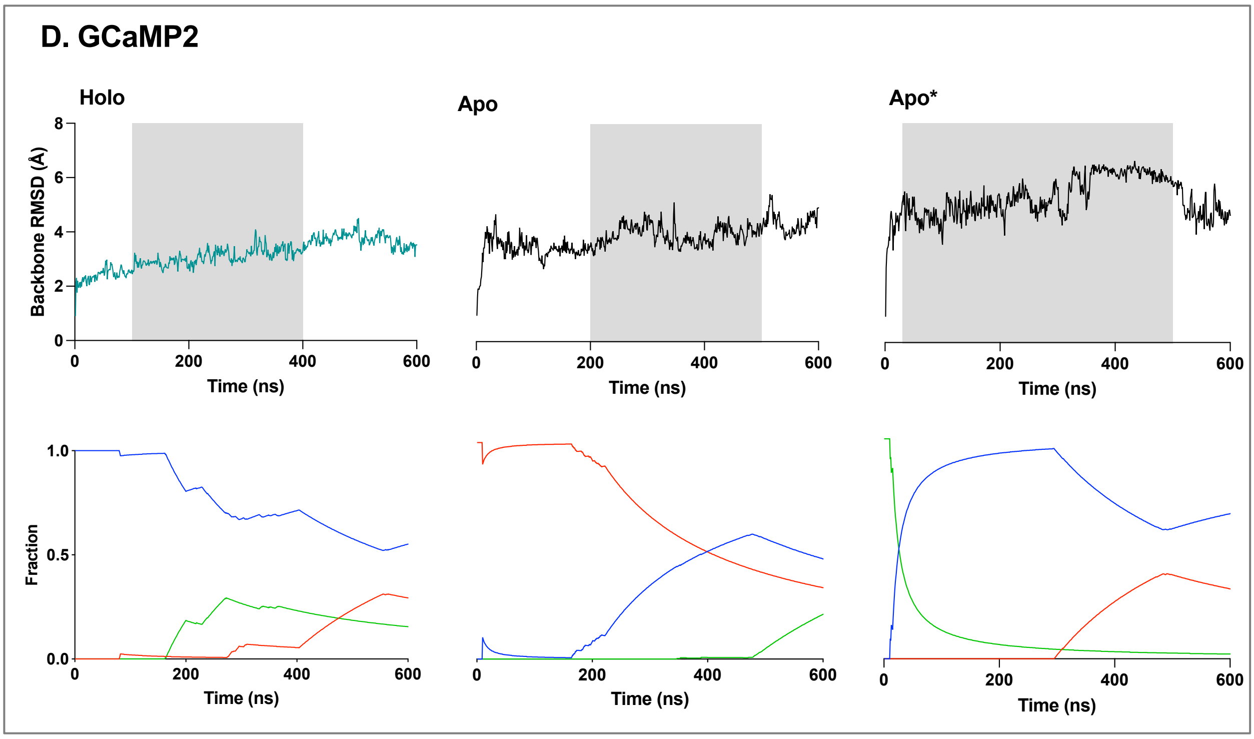


**Figure S2.** Backbone RMSD (upper) and conformational cluster fraction (lower) plots. Shaded area in RMSD plots indicates the part of the trajectory used in analysis. In cluster fraction vs time plots, each color represents a different a different conformational cluster.

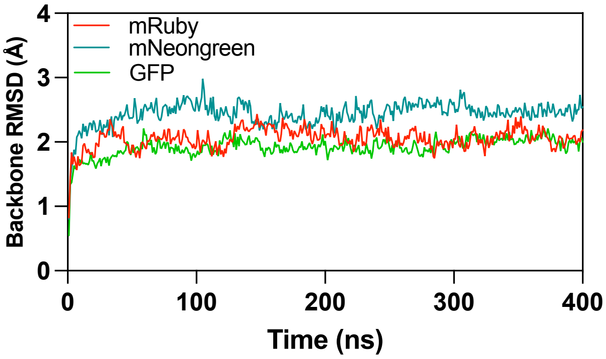


**Figure S3.** Backbone RMSD of ON state (anionic) parental FPs

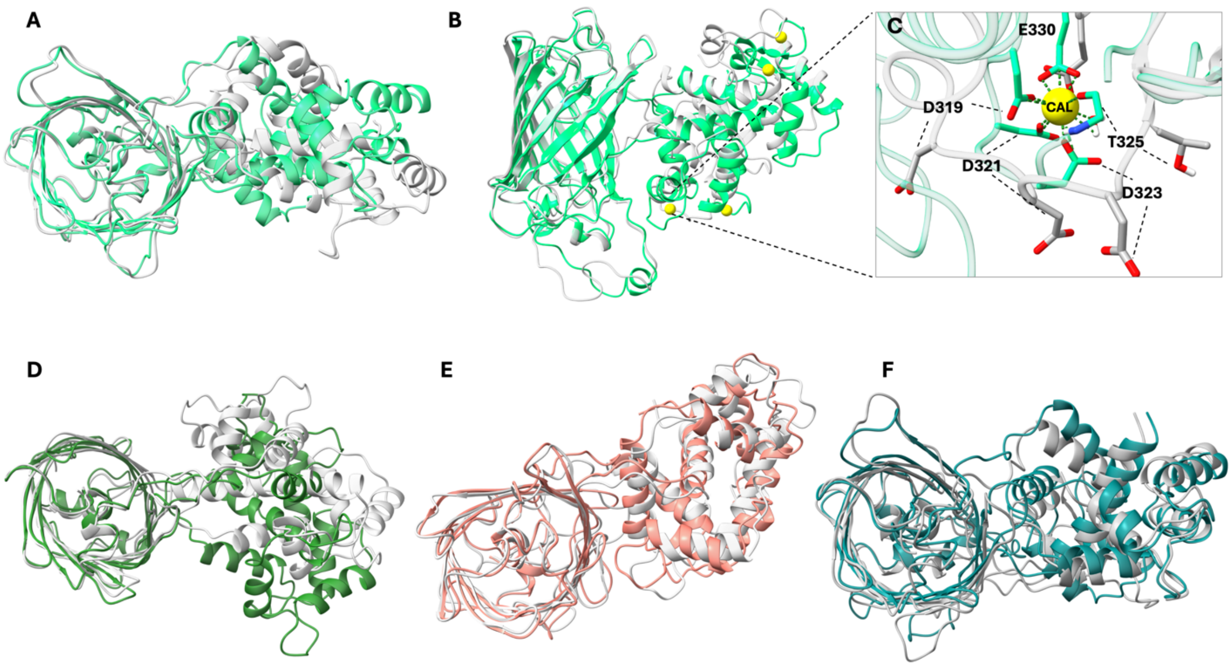


**Figure S4.** Changing positions of CaM and FP domains upon relative to each other as a result of calcium removal from the holo sensor. **A-C** jGCaMP8. Top **(A)** and side view **(B)** of apo and holo states aligned onto the FP domain. **C.** Calcium coordinating residues on EF2 hand. Top views of *holo* and *apo* states of **D.** NCaMP7 **E.** RCaMP **F.** GCaMP2. *Apo* states are colored silver in each case.

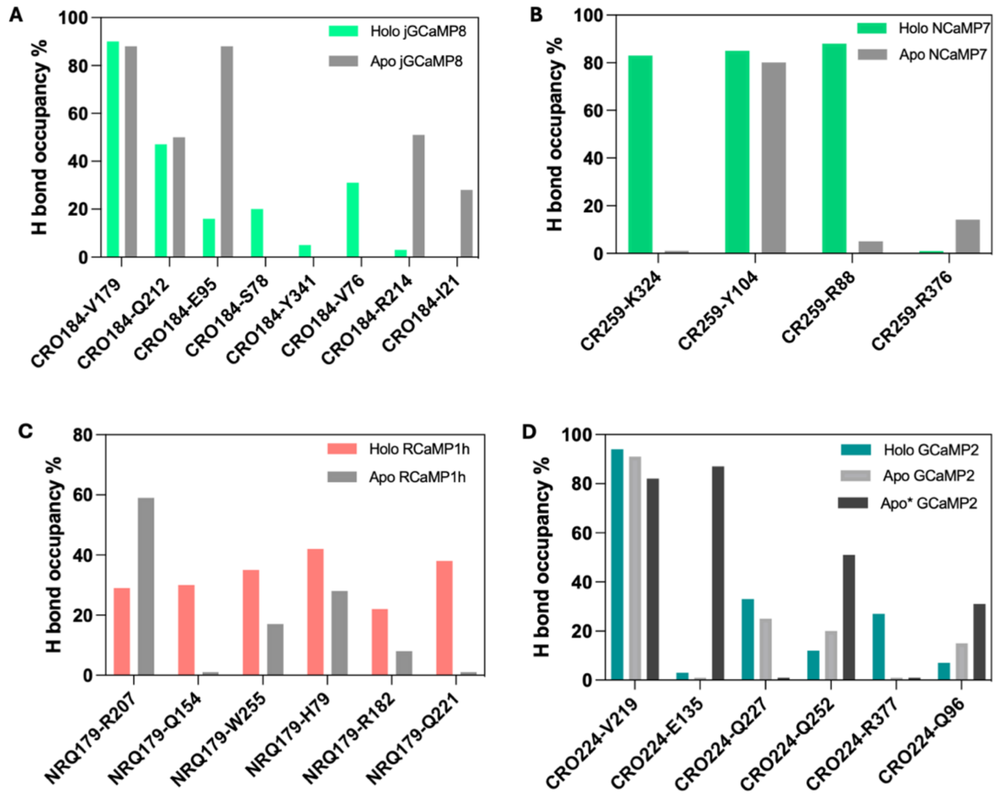


**Figure S5.** Occupancies of hydrogen bonds involving the chromophore as either acceptor or donor in holo and apo sensors. A. jGCaMP8 B. NCaMP7 C. RCaMP1h D. GCaMP2

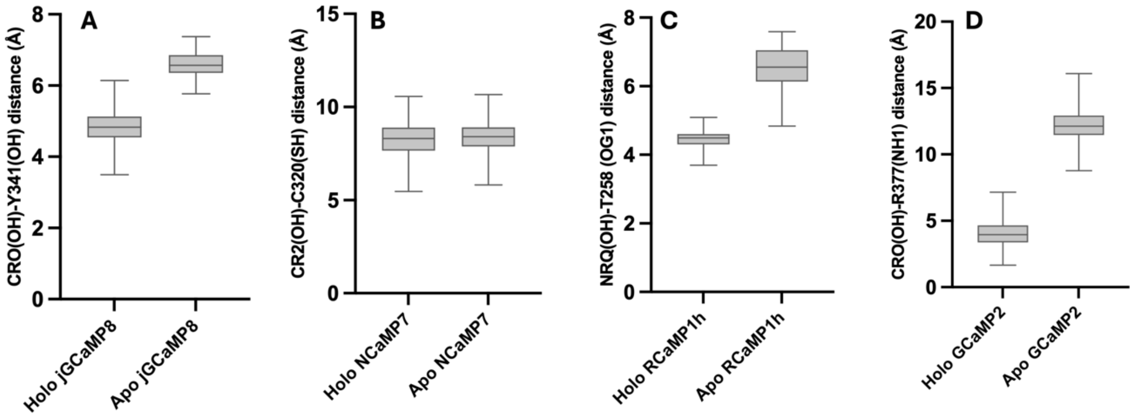


**Figure S6.** Distribution of interatomic distances between chromophore phenoxy oxygen and its primary hydrogen bond donor in the *holo* (ON) and *apo* (OFF) states. **A. jGCaMP8 B. NCaMP7 C. RCaMP1h D. GCaMP2**

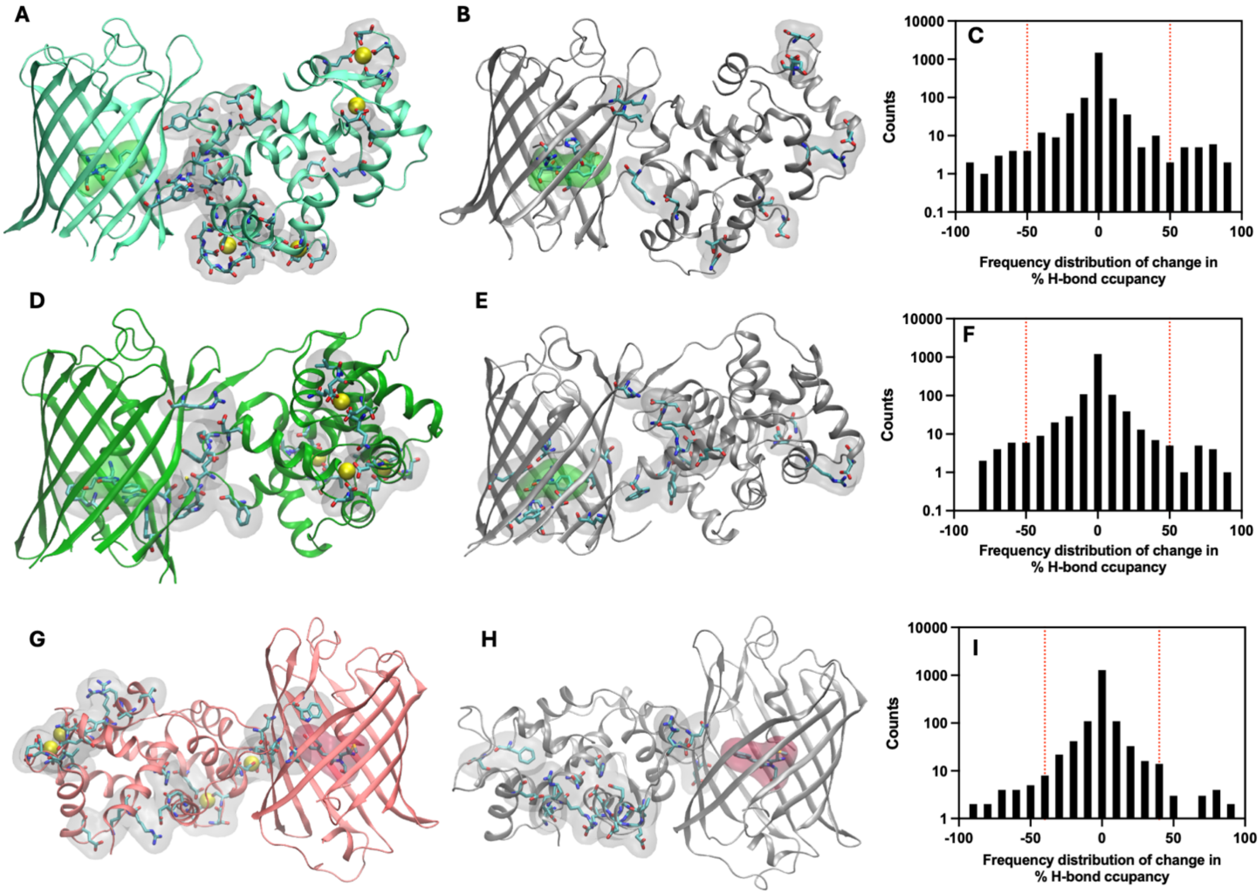


**Figure S7.** Shift in hydrogen bond occupancies between holo and apo sensors. **A-C** JGCaMP8 **D-E** NCaMP7 **G-I** RCaMP. *Apo* states are colored silver while *holo* states are colored based on the color emitted.

**Table S2.** Progressive change of hydrogen bond occupancies going from *holo* to *apo** state in GCaMP2

| *Residue pair* | *Location* | *Bond occupancy %* | | |
|---|---|---|---|---|
| | | *Holo* | *Apo* | *Apo** |
| R81-E387 | Interface | 96 | 7 | 0 |
| E370-A360 | CaM | 93 | 9 | 0 |
| K46-E314 | CaM | 65 | 7 | 0 |
| E317-K46 | CaM | 89 | 67 | 0 |
| D398-E407 | CaM | 86 | 18 | 0 |
| R42-E314 | CaM | 85 | 28 | 0 |
| T382-N83 | Interface | 79 | 42 | 0 |
| E417-K43 | CaM | 54 | 0 | 0 |
| T347-E350 | CaM | 80 | 1 | 21 |
| D323-F319 | CaM | 69 | 59 | 13 |
| R377-D381 | CaM | 62 | 0 | 0 |
| E357-N62 | Interface | 58 | 24 | 0 |

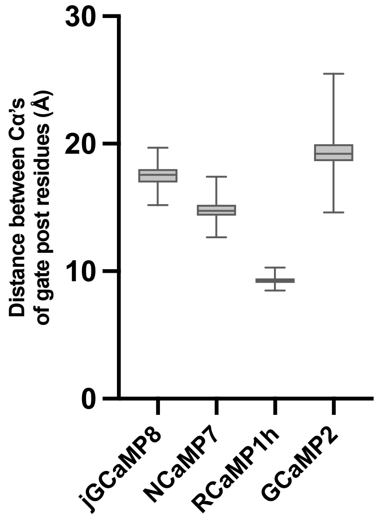


**Figure S8. Interatomic distances between $C_\alpha$ atoms of gate post residues in the holo state of each sensor.** jGCaMP8: I21-D65, NCaMP7: A315-C320, RCaMP1h: N25-W255, GCaMP2: E61-T203 distance

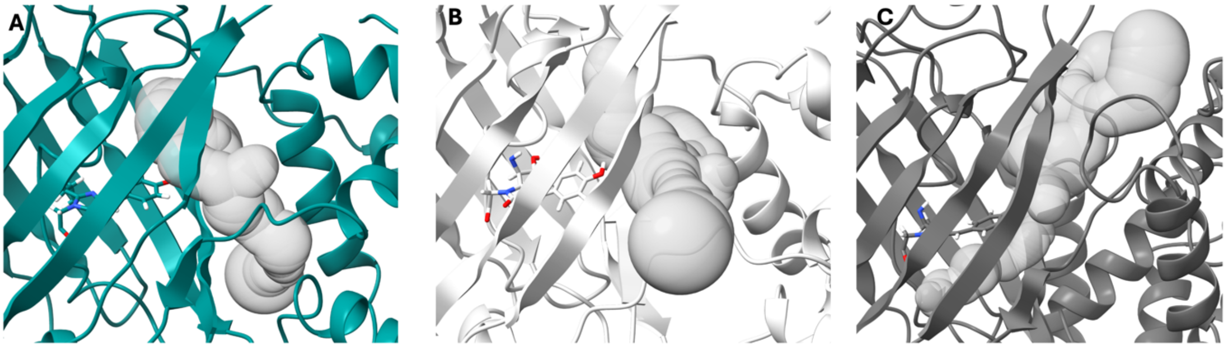


**Figure S9. Solvent channels of the various forms of GCaMP2. A.** GCaMP2 *holo* **B.** GCaMP2 *apo* **C.** GCaMP2 *apo** (calculated by Mole2.5)
